# *WEScover*: selection of whole exome sequencing vs. gene panel testing

**DOI:** 10.1101/367607

**Authors:** William Jefferson Alvarez, In-Hee Lee, Carles Hernandez-Ferrer, Kenneth D. Mandl, Sek Won Kong

## Abstract

**Motivation:** Whole exome sequencing (WES) is widely adopted in clinical and research settings. However, there is potential for false negatives due to incomplete breadth and depth of coverage for several exons in clinically implicated genes. In some cases, a targeted gene panel testing may be a dependable option to ascertain true negatives for genomic variants in known phenotype associated genes. We developed a tool for quickly gauging whether all genes of interest would be reliably covered by WES or whether targeted gene panel testing should instead be considered to minimize false negatives in candidate genes.

**Results:** *WEScover* is a novel web application that provides an interface for discovering breadth and depth of coverage across population scale WES datasets, searching either by phenotype, by targeted gene panels and by gene(s). Moreover, the application shows metrics from the Genome Aggregation Database to provide gene-centric view on breadth of coverage.

**Conclusion:** *WEScover* allows users to efficiently query genes and phenotype for exome coverage of associated exons, and recommends use of panel tests for genes that are potentially not well covered by WES.

## Background

As the cost of whole exome sequencing (WES) drops, WES is replacing broad and/or targeted gene panel testing [1, 2]. WES, for example, is superior in measurement of the ever-growing number of driver and passenger mutations in diverse genes across different cancer types as well as increasing awareness of polygenic contribution to most genetic disorders. However, WES may not capture all exons in clinically implicated genes in the human genome [3, 4] and whole genome sequencing (WGS) faces a similar challenge for some genes including highly polymorphic ones. Population scale aggregation of WES and WGS clearly shows limited breadth of coverage for some clinically implicated genes [3, 5]. Therefore, gene panel testing, whether for a single gene or for hundreds of candidate genes, is still a clinically useful measure where false negatives due to suboptimal coverage of WES and WGS are likely. Yet it is difficult to predict whether the exons known to harbor disease-associated variants would be covered with sufficient persite depth of coverage to reliably call variants or not.

*WEScover* provides the advantage of summarizing coverage information on clinically implicated genes and highlighting population-specific differences in exome coverage. This summary can provide a basis to recommended the use of gene panel tests for the genes that are poorly covered by WES. Also, *WEScover* provides WES coverage stratified by continental-level population groups. With a self-reported ancestry of the patient,users would find the coverage of a given gene that matches the population-specific coverage compared to other datasets such as the Genome Aggregation Database project [6] that only provides global mean coverage across all exomes. Links to gnomAD are provided such that continental-level and global coverage metrics could be compared.

## Implementation

To help biomedical investigators to select the reliable genetic testing strategy – i.e., WES vs. targeted gene panel(s), we developed the *WEScover*, web application that highlights global gene level coverage and inter-individual variation in breadth of coverage for genes along with corresponding genetic tests listed in the National Institutes of Health Genetic Testing Registry (GTR) [7]. A total of 6,097 putative disease-associated genes are listed across 46,104 genetic tests for both clinical and research usage including 32,275 CLIA-certified ones in GTR (last access: 2019-12-13, updated daily). For each unique exon in the Consensus Coding Sequence (CCDS) [8], we calculated breadth of coverage at >10x, >20x and >30x (the percentage of sites where per-site depth of coverage is higher than 10x, 20x, and 30x, respectively) across the exomes from the 1000 Genomes Project (1KGP) [9] phase 3 (N=2,504, alignment files remapped to GRCh38 human reference genome). Additionally, we took the average value of the full exomes (N=123,136) from gNOMAD as global estimates from a large-scale data (the continent-level data is not currently available in the gnomAD project). Using the relationship between phenotypes, listed either in GTR or Human Phenotype Ontology (HPO) [10], genetic test names from GTR, and genes, we created a database and a query interface as a R Shiny application (package version 1.3.2) [11].

## Results

The initial query interface allows users to enter phenotype, genetic test name (retrieved from the GTR website), or official gene symbol(s) of interest. For each gene matching the query, the global mean of breadth of coverage along with its maximum and minimum values is shown as a table in an ascending order of global means (**Figure 1A**). By default, we used breadth of coverage at > 20x – a threshold sufficient to achieve 99% sensitivity for detecting single nucleotide variant [12]. We also performed a one-way analysis of variance to test differences between means of populations and reported the test statistics and p-value in this table. The button at the end of each row opens a panel with further details about the coverage of the gene. The panel first shows a table with the mean of breadth of coverage stratified by continent-level population. The second tab shows a violin plot for breadth of coverage stratified by continent-level populations with the mean value from exomes in gnomAD project as a black line (**Figure 1B**). A plot for coverage at each genomic position of the selected gene, based on gnomAD coverage data, is shown next to the violin plot (Fig. 1C). Lastly, the panel reports all genetic test involving the gene. Insufficient breadth of coverage in both projects, 1KGP and gnomAD, should warn the user that the candidate genes may not be well covered in WES and that targeted gene panel tests should be considered to minimize potential false negatives.

**Figure 1.**
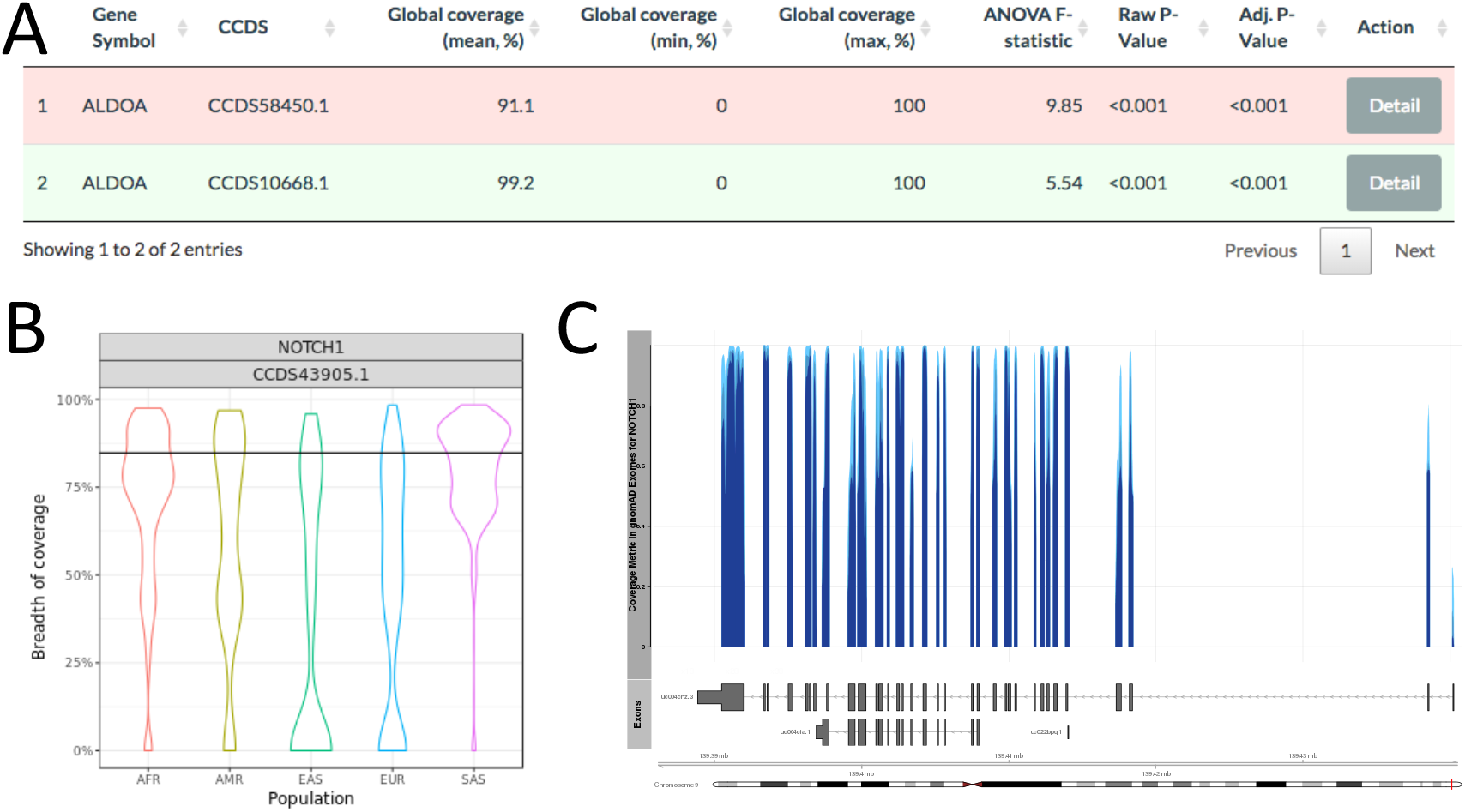
A) The initial screen for selected genes. Clicking ‘Detail’ button (red highlighted box) shows a window with more information for the selected transcript such as breadth of coverage per contintent-level population (B), coverage metric from gnomAD Exomes (C), and list of gene panels including the transcript for selected transcript. B) The violin plot shows the distribution of coverage metrics from 1KGP exomes in each of five continent-level population. The black horizontal line denotes the global average value from gnomAD exomes. C) The coverage plot shows the transcript model and coverage metric from gnomAD exomes. The upper part of graph shows metric values at 10x (most light blue), 20x, and 30x (most dark blue).

## Conclusion

WES and WGS provide comprehensive evaluation of genomic variants in various conditions; however, users must be informed regarding possible false negatives due to incomplete breadth and depth of coverage (ideally, from sequencing vendors). In such cases, targeted gene panel tests should be considered as a primary choice over the others. *WEScover* can guide users as to whether WES is appropriate for testing the genes of interest. Together with information from GTR, which provides transparent and comprehensive list of genetic tests with indications, users can make an informed decision for testing genes prior to ordering genetic tests in the clinical settings.

## Availability and requirements

**Project name**: *WEScover*

**Project home page**: https://tom.tch.harvard.edu/shinyapps/WEScover/

**Project source code**: https://github.com/bch-gnome/WEScover

**Operating system**: Platform independent

**Programming language**: R Shiny

**Other requirements**: *WEScover* requires the following R packages: *shiny*, *shinythemes*, *DT*, *ggplot2*, *shinyjs*, *reshape2*, *RColorBrewer*, *fst*, *data.table*, and *corrplot*.

**License**: MIT

**Any restrictions to use by non-academics**: None

## Abbreviations

WES: Whole exome sequencing
WGS: Whole genome sequencing
gnomAD: Genome Aggregation Database
GTR: Genetic Testing Registry
CCDS: Consensus Coding Sequence
1KGP: 1000 Genomes Project
HPO: Human Phenotype Ontology

## Declarations

### Ethics approval and consent to participate

Not applicable.

### Consent for publication

Not applicable.

### Availability of data and materials

Breadth of coverage data by continent-level populations from 1000 Genomes Project and global exome coverage plots in gnomAD are available for download in https://tom.tch.harvard.edu/shinyapps/WEScover/ under the ‘Data’ tab.

### Competing interests

The authors declare that they have no competing interests.

### Funding

This work has been supported by the Boston Children’s Hospital Precision Link initiative. SWK was supported in part by grants from the National Institutes of Health (R01MH107205, R24OD024622, U01TR002623 and U01HG007530).

### Authors’ contributions

IHL and SWK generated the original breadth of coverage data summarized in *WEScover*. The source code of the web interfae application was developed by WJA, IHL, and CHF. SWK drafted the initial manuscript; KDM provided funding and critical evaluation of the manuscript. All authors have read and approved the final manuscript.

## Acknowledgments

Not applicable.

